# Household and climate factors influence *Aedes aegypti* risk in the arid city of Huaquillas, Ecuador

**DOI:** 10.1101/2020.05.19.104372

**Authors:** James L. Martin, Anna M. Stewart-Ibarra, Efraín Beltrán Ayala, Erin A. Mordecai, Rachel Sippy, Froilán Heras Heras, Jason K. Blackburn, Sadie J. Ryan

**Affiliations:** Quantitative Disease Ecology and Conservation (QDEC) Lab, Department of Geography, University of Florida, Gainesville, Florida, United States of America; Emerging Pathogens Institute, University of Florida, Gainesville, Florida, United States of America; Institute for Global Health & Translational Sciences, SUNY Upstate Medical University, Syracuse, New York, United States of America; Department of Medicine, SUNY Upstate Medical University, Syracuse, New York, United States of America; InterAmerican Institute for Global Change Research (IAI), Montevideo, Uruguay; Universidad Técnica de Machala, Machala, Ecuador; Biology Department, Stanford University, Stanford, California, United States of America; Spatial Epidemiology and Ecology Research Laboratory, Department of Geography, University of Florida, Gainesville, Florida, United States of America

## Abstract

Arboviruses transmitted by *Aedes aegypti* (e.g., dengue, chikungunya, Zika) are of major public health concern on the arid coastal border of Ecuador and Peru. This high transit border is a critical disease surveillance site due to human movement-associated risk of transmission. Local level studies are thus integral to capturing the dynamics and distribution of vector populations and social-ecological drivers of risk, to inform targeted public health interventions. Our study examines factors associated with household-level *Ae. aegypti* presence in Huaquillas, Ecuador, while accounting for spatial and temporal effects. From January to May of 2017, adult mosquitoes were collected from a cohort of households (n = 63) in clusters (n = 10), across the city of Huaquillas, using aspirator backpacks. Household surveys describing housing conditions, demographics, economics, travel, disease prevention, and city services were conducted by local enumerators. This study was conducted during the normal arbovirus transmission season (January - May), but during an exceptionally dry year. Household level *Ae. aegypti* presence peaked in February, and counts were highest in weeks with high temperatures and a week after increased rainfall. Presence of *Ae. aegypti* was highly variable between clusters. Hierarchical generalized linear models were used to explore household social-ecological variables and female *Ae. aegypti* presence. Houses with *Ae. aegypti* used larvicide in water tanks and had high awareness of dengue transmission. We found that homes were more likely to have *Ae. aegypti* when heads of household had lived in the neighborhoods for longer than average (>22 years), when households had more occupants than average (>4.5), had a female head of household, and received more frequent garbage collection. *Ae. aegypti* presence was less likely in households with reliable water supply and septic systems. Based on our findings, infrastructure access, urban occupancy patterns, and seasonal climate are important considerations for vector control in this city, and even in dry years, this arid environment supports *Ae. aegypti* breeding habitat.

**Author summary:** Mosquito transmitted infectious diseases are a growing concern around the world. The yellow fever mosquito (*Aedes aegypti*) has been responsible for recent major outbreaks of disease, including dengue fever and Zika. This mosquito prefers to bite humans and lay its eggs in artificial containers such as water tanks and planters. This makes *Ae. aegypti* well suited to become established in growing urban areas. Controlling these mosquitoes has been an important way to reduce the risk of disease transmission. Studies that are undertaken to understand local factors that contribute to the continued survival of the mosquito can be used to inform control practices. We conducted a study in the largest city on the border of Ecuador and Peru where we collected adult mosquitoes from houses and surveyed household members about their behaviors, perceptions, and housing infrastructure associated with the mosquito vector. Mosquitoes were most numerous in weeks with high temperatures and a week after increased rainfall. Larvicide was a commonly used control strategy in homes where *Ae. aegypti* was present. We found that houses that had more people, female heads of household, heads of household that had lived in the neighborhood for a long time, and had unreliable water service, were more likely have mosquitoes present, while houses that used septic systems were less likely to have mosquitoes present.

## Introduction

Arboviral diseases are an increasing global concern [1]; dengue fever has the largest health burden, with 58.4 million cases reported annually worldwide [2]. The *Aedes aegypti* (*Ae. aegypti*) mosquito is the primary vector of dengue virus and other important arboviruses such as chikungunya, yellow fever, and Zika [1,3]. This mosquito is well-adapted to urban environments as it is an anthropophilic container breeder [4]. Increasing urbanization and trends in international trade and travel have greatly facilitated the spread and establishment of *Ae. aegypti* and the diseases it transmits [5]. Vector control remains the primary strategy to prevent arboviral diseases [6], and it has become increasingly apparent that the social and ecological factors that influence vector populations vary greatly in space and time [7,8]. This has significant implications for the effectiveness of vector control programs. Local studies are therefore necessary to assess place-specific drivers and to inform interventions [9]. *Ae. aegypti* mosquitoes are sensitive to abiotic factors such as climate, including temperature, rainfall and relative humidity [10]. Its anthropophilic nature also makes the mosquito sensitive to human activities [11] such as household water storage or use of repellant. Thus, it is important to examine both the social and ecological components of a local environment to understand subcity risks of vector presence, and the potential for intervention.

International border regions represent an interesting opportunity for the study of disease vectors. Neighboring countries may be a source of infected vectors or imported human cases of disease [12]. Differences in environment, socio-economic status, access to healthcare, healthcare practices, and vector control practices between countries can impact the burden of disease, abundance of vectors, and infection levels of vectors [12,13]. Historically, international transportation of goods and people has introduced or re-introduced *Ae. aegypti, Aedes albopictus* (a secondary vector of dengue) and dengue virus to new locations [14–16]. Cities along borders may also have high levels of migrant populations or may be a through-way for migrating populations, who can serve as reservoirs of vector-borne disease [17,18].

Ecuador has a history of high arboviral disease burden [19,20]. Yellow fever dominated during early 20th century, and dengue fever emerged as the principal mosquito-borne disease at the beginning of the 21st century with the decline in malaria cases [21,22]. More recently, the introduction of new viruses, such as chikungunya and Zika, has resulted in major epidemics [23]. There have been multiple studies in recent years of climate, social-ecological factors, and *Ae. aegypti* in Ecuador [11,19,24–29]. These neighborhood-level studies have primarily focused on Machala, Ecuador, an urban center with a steppe climate (BSh) per the Köppen climate classification system (e.g., intermediate between desert and humid climate). Studies found that both climate and social-ecological factors influenced the presence of immature and adult *Ae. aegypti* in neighborhoods [11,24–29]. The El Niño–Southern Oscillation is an important driver of the climates of southern Ecuador and northern Peru [27]. To date, there have been no studies of the impacts of climate and social-ecological system (SES) on *Ae. aegypti* in the desert climates of Ecuador, nor along border regions.

This study aims to identify the climatic and social-ecological aspects of household-level *Ae. aegypti* presence in the arid border city of Huaquillas. These insights can lead to a better understanding of how this vector-borne disease system functions and where potential levers of control might be found. The knowledge gained can help inform intervention decisions in Huaquillas and other similar settings. Additionally, this study, in combination with similar local-scale studies in the region, can help answer questions relating to scale and heterogeneity in arboviral disease systems.

## Methods

### Ethical review

This study protocol was reviewed and approved by Institutional Review Boards (IRBs) at SUNY Upstate University, the Luis Vernaza hospital in Guayaquil, Ecuador, and the Ministry of Health of Ecuador. Prior to the start of the study, adult participants (≥ 18 years of age) engaged in a written informed consent (conducted in Spanish). To protect the privacy of participants, data were de-identified prior to use in any of the analyses conducted in this study.

### Study site

Huaquillas is a coastal city in southern Ecuador with a population of 48,285 (Fig 1) [30]. It is situated on the border of Ecuador and Peru and is the primary crossing at the southern border, and a major hub of transit between the two countries [31]. In terms of total migration, Huaquillas is the third most active city in Ecuador, with 622,405 arrivals and departures annually [31]. There is frequent economic exchange with the similarly-sized Peruvian city of Zarumilla located less than 2 km from the border [32,33]. Increased binational cooperation and a relaxation of trade and travel barriers in recent decades has contributed to increased urban development on both sides of the border [33].

**Fig 1.**
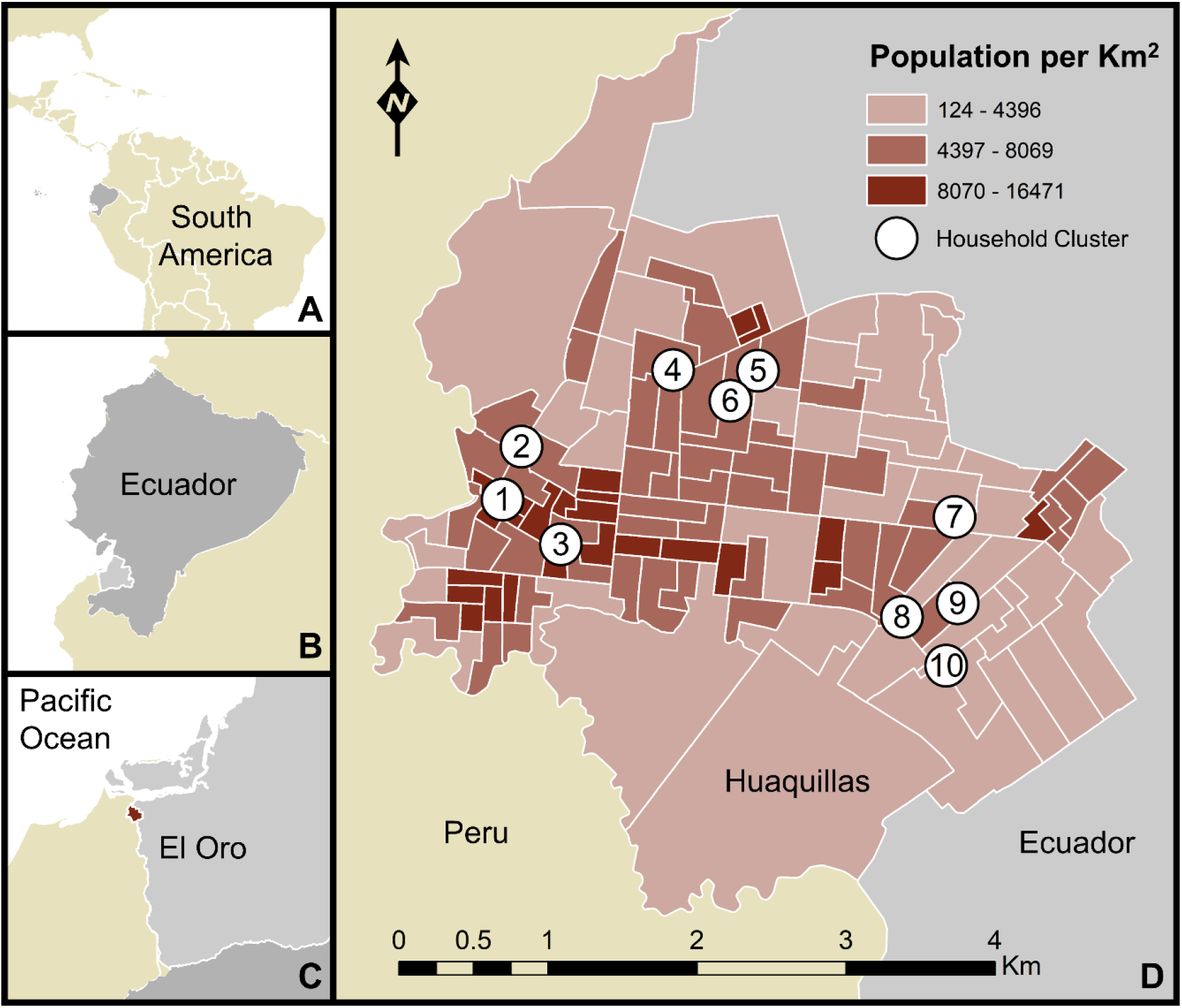
Map of the study area. Huaquillas is located within South America, Ecuador, and El Oro province. The map includes household cluster locations where sampling took place (white circles denote areas where up to 5 houses were sampled, but precise household locations are not shown, to protect identities) and the population density of Huaquillas at the census tract level. a) location of Ecuador, b) location of El Oro province, c) location of Huaquillas (red) d) Huaquillas population and sampling sites.

Huaquillas is in a low-lying coastal region highly suitable for *Ae. aegypti*, as underscored by major outbreaks of dengue fever in recent years [27]. It has a hot desert climate (BWh) per the the Köppen climate classification system [34]. While desert climates are often characterized by limited precipitation, intense sunlight, and little vegetation, actual conditions can vary greatly by place. Huaquillas experiences a monsoon season which occurs during the first half of the year, with monthly total rainfall peaking in February at 128 mm (Fig 2). Increased precipitation coincides with high temperatures. Annual mean temperature ranges from 21.6-27.6°C. Minimum daily temperatures range from 18.8—22.3°C annually and maximum daily temperatures range from 25.8—33.0°C annually.

**Fig 2.**
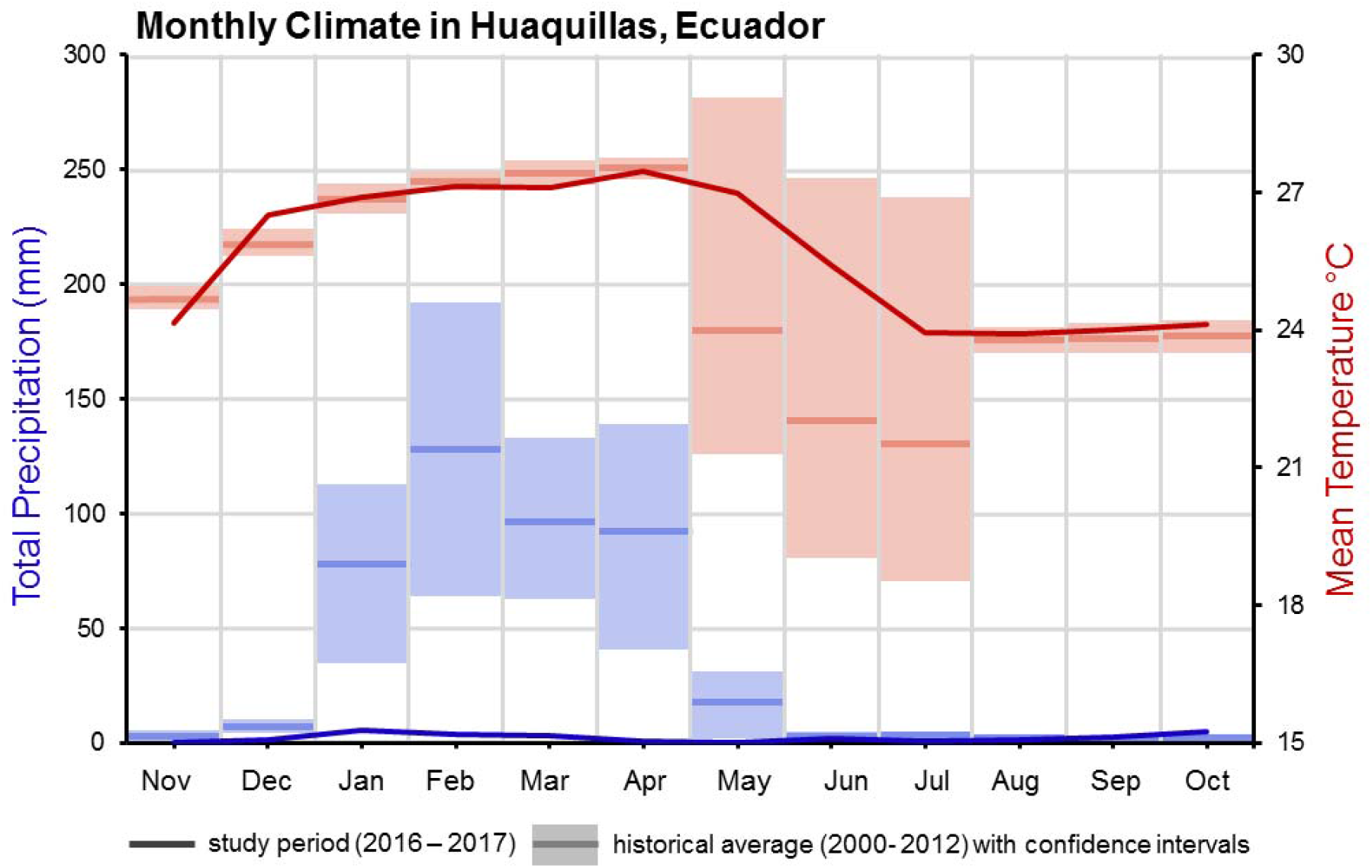
Climate in Huaquillas, Ecuador. Monthly mean temperature in red and total monthly precipitation as blue. Solid lines represent the climate during the study period (2016-2017) while box plots represent the climatology from 2000-2012 (historical monthly averages and 95% confidence intervals).

### Data sources

#### Climate data

The National Institute of Meteorology and Hydrology (INAMHI) provided daily climate data for Huaquillas from 2000—2012 and hourly data (March 2016—December 2017) from an automatic weather station in the city. Both datasets include minimum, maximum, and mean temperature, and total daily precipitation.

#### Census data

Tract-level census data from the most recent national census (2010) was provided by the National Institute of Statistics and Census (INEC) and were assigned to georeferenced tract polygons were provided by INAMHI.

#### Participating households

Households were selected for inclusion in this study using a semi-random selection process (cluster design). The study was designed to capture clusters of up to five households facing similar risks within the radius of the flight range of an *Ae. aegypti* mosquito. Each cluster included a central household with four or five nearby households, preferably within a 250 m radius to represent the flight distance of *Ae. aegypti* [35]. 10 clusters were located across the city. The resulting sampling design was a favorable balance between sufficiently accounting for the heterogeneity of the Huaquillas urban environment and the logistical feasibility of regularly visiting the households. Houses were georeferenced and given specific house and cluster codes.

#### Household mosquito samples

Adult mosquitoes were collected from households biweekly during January—May 2017, using backpack aspirators. Not all clusters were sampled in each sampling event, for logistical reasons. Adult mosquitoes were stored on ice in a cooler and were transported to the entomology lab at the Universidad Técnica de Machala, Ecuador, 74km by vehicle. Specimens were enumerated and sorted by sex and species (*Ae. aegypti* and other).

#### Household surveys

The head of each household responded to a survey about household demographics, occupation, access to public services, knowledge and perceptions of mosquito-borne disease, mosquito control and prevention, and household expenditures [36]. Study personnel completed a visual assessment of housing conditions, as in previous studies in the region [11,36]. Together, these responses comprise the social-ecological system (SES) variables available for analyses.

### Data Analyses

All data processing and analyses were conducted in R version 3.5.1 [37] using the packages “rgdal” [38], “betareg” [39,40], “lrtest” [41], “lme4” [42], “glmulti” [43] and “MuMIn” [44].

#### Climate data

Daily climate data from 2000-2012 were aggregated to month by summing precipitation readings and averaging minimum, maximum, and mean temperatures. A long-term monthly mean of each variable was calculated and 95% confidence intervals were constructed using an ordinary bootstrap procedure with 1000 replicates [45,46]. These long-term data served as the baseline for examining the monthly climate of Huaquillas in 2017.

Hourly climate data from 2016 and 2017 were aggregated to weekly measures. Hourly precipitation readings during that period were summed for weekly precipitation. Hourly temperature data were used to derive daily mean, minimum, and maximum temperatures.

#### Household mosquito samples

Household *Ae. aegypti* counts were summarized as presence: sampling events were classified as “positive” if female *Ae. aegypti* mosquitoes were present, and the proportion of positive households was calculated for each week. Household mosquito sampling occurred at 2 or 3 week intervals, so *Aedes aegypti* presence data were discontinuous [47]. Cross correlations using Pearson’s r were calculated for *Ae. aegypti* presence and climate variables (precipitation, mean, maximum, and minimum temperature) lagged from 0 to 6 weeks, with statistical significance assessed using t-tests. Statistically significant climate indicators were used to construct a beta regression model to capture the relationship between climate and positive households, and multi-model selection used to derive the best-fit model of these climate factors, using the package “glmulti” in R, and “MuMin” to create a weighted model average between models falling within 2 AICc of the minimum (top model).

#### Assessing temporal and spatial signals

To assess whether there was a temporal signal in household-level presence of female *Ae. aegypti* in the study group, at a monthly scale, only households with sampling events in all months were included. For each household, if multiple collections occurred in a month, one collection record was randomly selected to represent *Ae. aegypti* presence in that month.

To assess whether there was a spatial signal among the clusters of households in the study, for months and then clusters, the proportion of households with *Ae. aegypti* was calculated, and 95% binomial confidence intervals were derived using the Wilson score interval [48]. Pearson’s chi-squared tests were used to assess differences in *Ae. aegypti* presence between months and between clusters. Fisher’s exact test was used as a post hoc analysis to assess which groups were significantly different from one another. To reduce the chance of observing spurious differences between groups, which often occurs when conducting multiple comparisons on a set of data, p-values were adjusted using the false discovery rate method [49].

#### Bivariate tests for social factors

Associations between survey responses and household female *Ae. aegypti* presence during the study period were measured using bivariate tests. Questions that addressed social-ecological factors hypothesized to be important for vector population dynamics at this study site (based on previous studies in this region) were selected from the full household survey for analysis [11,25,36,50]. Hypothesized factors included water storage practices, building materials, and economic status. Questions that had a low rate of response were excluded to minimize observations discarded due to missing data. Questions that had the same response for nearly every observation were also excluded because they offer little ability to differentiate between houses with and without *Ae. aegypti*. Bivariate statistical tests were used assess to differences in survey responses by *Ae. aegypti* presence (Fisher’s exact test for binary responses and t-tests for numerical responses). These tests did not consider the month of each mosquito sample event.

#### Social-ecological system (SES) models of household factors and Aedes aegypti presence

Data inclusion criteria for the SES models were the most stringent. Households were required to have mosquito samples in all months; those with more than one sampling event per month had one event randomly selected for inclusion.

Using an information-theoretical approach, competing models of household female *Ae. aegypti* presence as a function of SES variables found to be significant in bivariate analyses were constructed using the ‘dredge’ function in package “MuMin” in R. Using Akaike’s Information Criterion corrected for small sample size (AICc) [51], an exhaustive search of all possible combinations of SES variables was conducted. The top model had the lowest AICc score, and competing models were within two AICc points of the top model. The relative importance of each variable in the top and competing models was assessed using summed Akaike weights, which considers both the frequency a variable appears in the models, as well as the performance of these models [44,52]. All models included random effects of cluster and month to account for the unobserved differences between groups of observations [53]. We created a weighted average model of all models within 2 AICc points of the top model.

## Results

Sixty-three households participated in the study. Fifty-eight heads of household (HOHs) (92%) responded to the survey. The average distance between households within clusters was 86 m. A total of 458 mosquito collections occurred over 10 individual weeks of sampling. For assessments of spatial and temporal signals, 41 houses (65%) with 205 mosquito sampling events met the criteria for inclusion. For assessments of household characteristics, 43 houses (68%) with 189 mosquito sampling events met the criteria for inclusion. For SES models, 32 houses (51%) with 160 mosquito sampling events met the criteria for inclusion.

### Climate

Weekly *Ae. aegypti* female presence was significantly correlated with precipitation, with the strongest correlation at 1-week lag (r= 0.76, p = 0.001), and another significant lag at 3 weeks (r=0.66, p=0.036). Weekly *Ae. aegypti* female presence was also significantly correlated with mean (r=0.75, p=0.012) and maximum (r=0.84, p=0.002) temperature, in the same week. Minimum temperatures were not significantly correlated with *Ae. aegypti* presence at any lag.

A multi-model selection on a beta regression of the proportion of households with female *Ae. aegypti* presence as a function of mean temperature, maximum temperature, and precipitation for the week, and precipitation lagged 3 weeks, yielded two models within 2 AICc of the top model, and an averaged model of *Ae. aegypti* presence as a function of mean temperature, and precipitation lagged 1 week, had a pseudo R^2^ of 0.71. The current week’s temperature was significant (*β* = 0.63, SE=0.29, p=0.03) in the overall model, while precipitation of the prior week was not (*β* = 0.74, SE= 1.94, p=0.051).

### Temporal and spatial patterns of Ae. aegypti presence

The proportion of positive households was highest in February and lowest in May (Fig 3). In January, the proportion of positive households was 0.38; this increased to 0.59 in February before steadily declining. The monthly differences were significant (χ^2^=9.564, p=0.048), but a post hoc Fisher’s exact test did not identify specific month-to-month differences (S1 Table).

**Fig 3.**
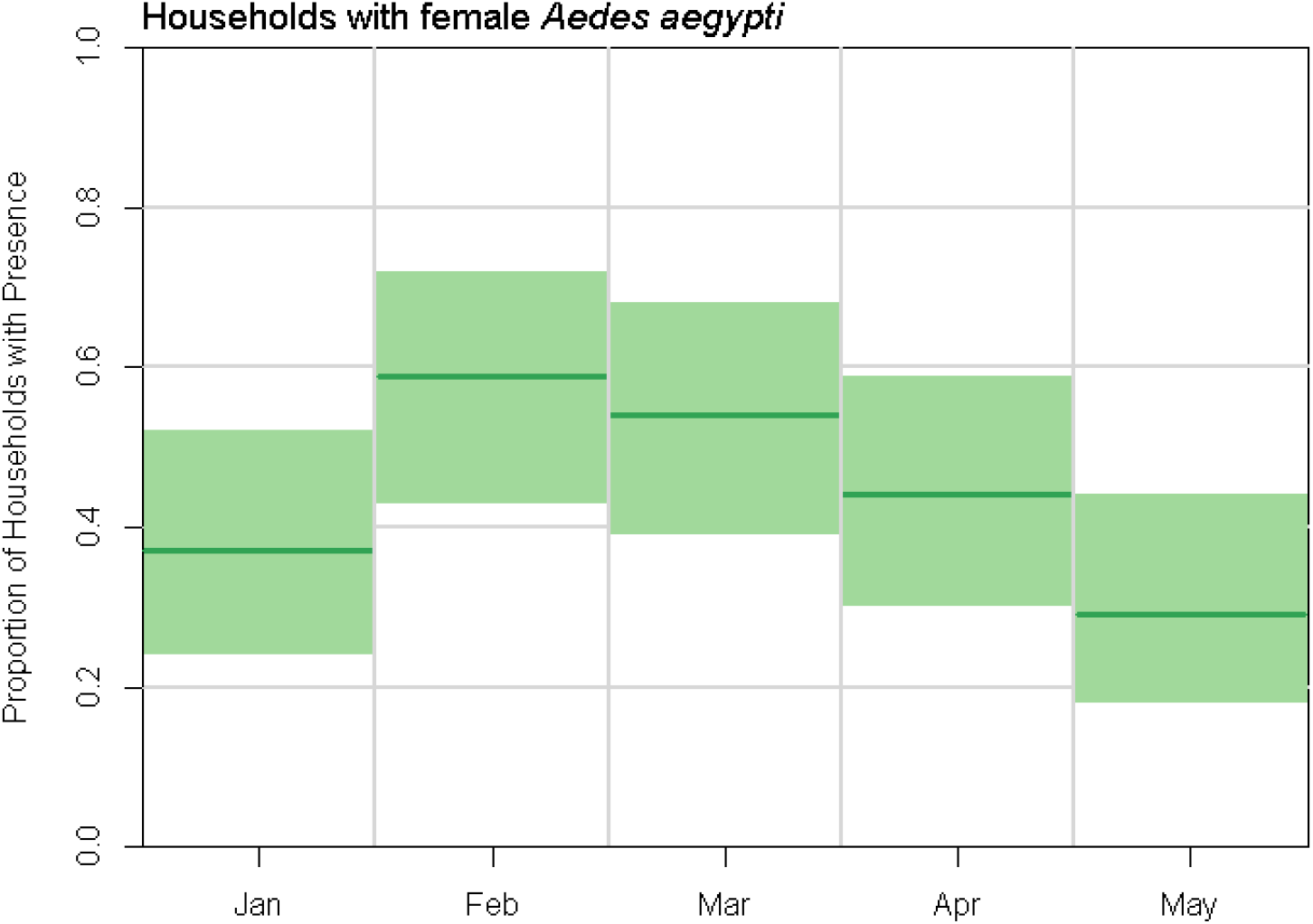
Female *Aedes aegypti* presence in households in Huaquillas, Ecuador, by month in 2017. Month proportions and 95% confidence intervals for 41 households used in this dataset. Chi-squared test for difference in proportions: 9.56, p=0.0484).

The positive proportion of houses by cluster varied from 0.15 to 0.70 (S2 Table). There were significant differences between clusters (χ^2^=32.185, p=0.0002). A post hoc Fisher’s exact test identified significant pairwise differences between four cluster pairs (Fig 4 and S3 Table).

**Fig 4.**
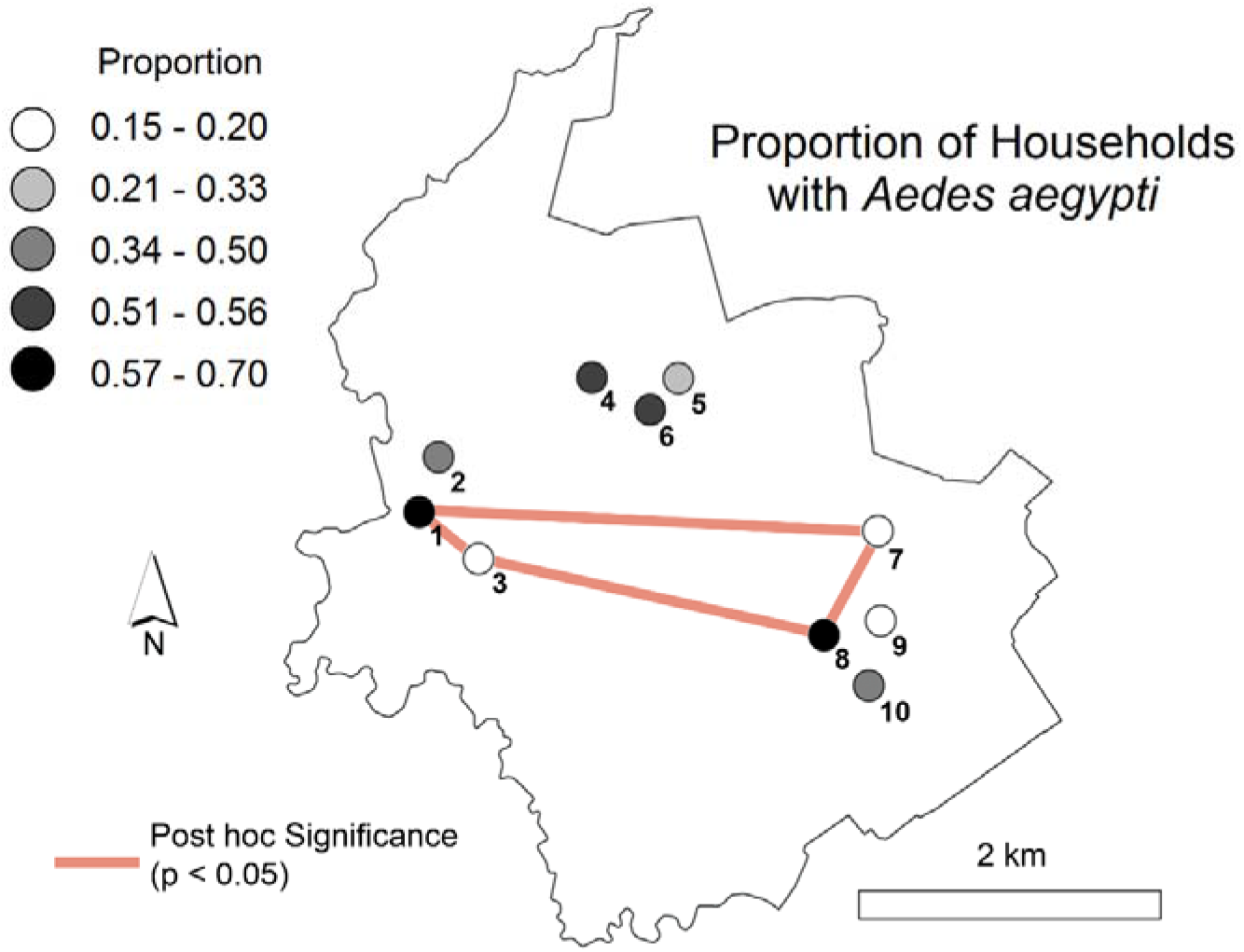
Proportion of households with female *Aedes aegypti* by cluster. The proportion of sampling events with female *Aedes aegypti* mosquitoes during January-May is indicated by shading (grey). Orange lines connecting clusters indicate a statistically significant difference in *Aedes aegypti* presence (post-hoc Fisher’s exact test).

### Bivariate tests for SES factors

All respondents considered dengue fever to be a severe disease and correctly answered questions about the transmission cycle of the disease, so these variables were not included in this analysis. Among demographic variables, the number of people living in a household, the number of years a head of household (HOH) had lived in the neighborhood, and female HOHs were significantly associated with household presence of female *Ae. aegypti* (Table 1). Factors describing the physical conditions of households were not found to be significant predictors of *Ae. aegypti* presence. However, three of the four variables describing infrastructure and public services were significant. Experiencing frequent interruptions in piped water service and biweekly trash collection were positively associated with presence, while use of a septic tank was negatively associated. Of mosquito prevention methods surveyed, only reported use of Abate^®^ larvicide, was significant, with a positive effect. Employment and travel habits were not found to be significant.

**Table 1.**
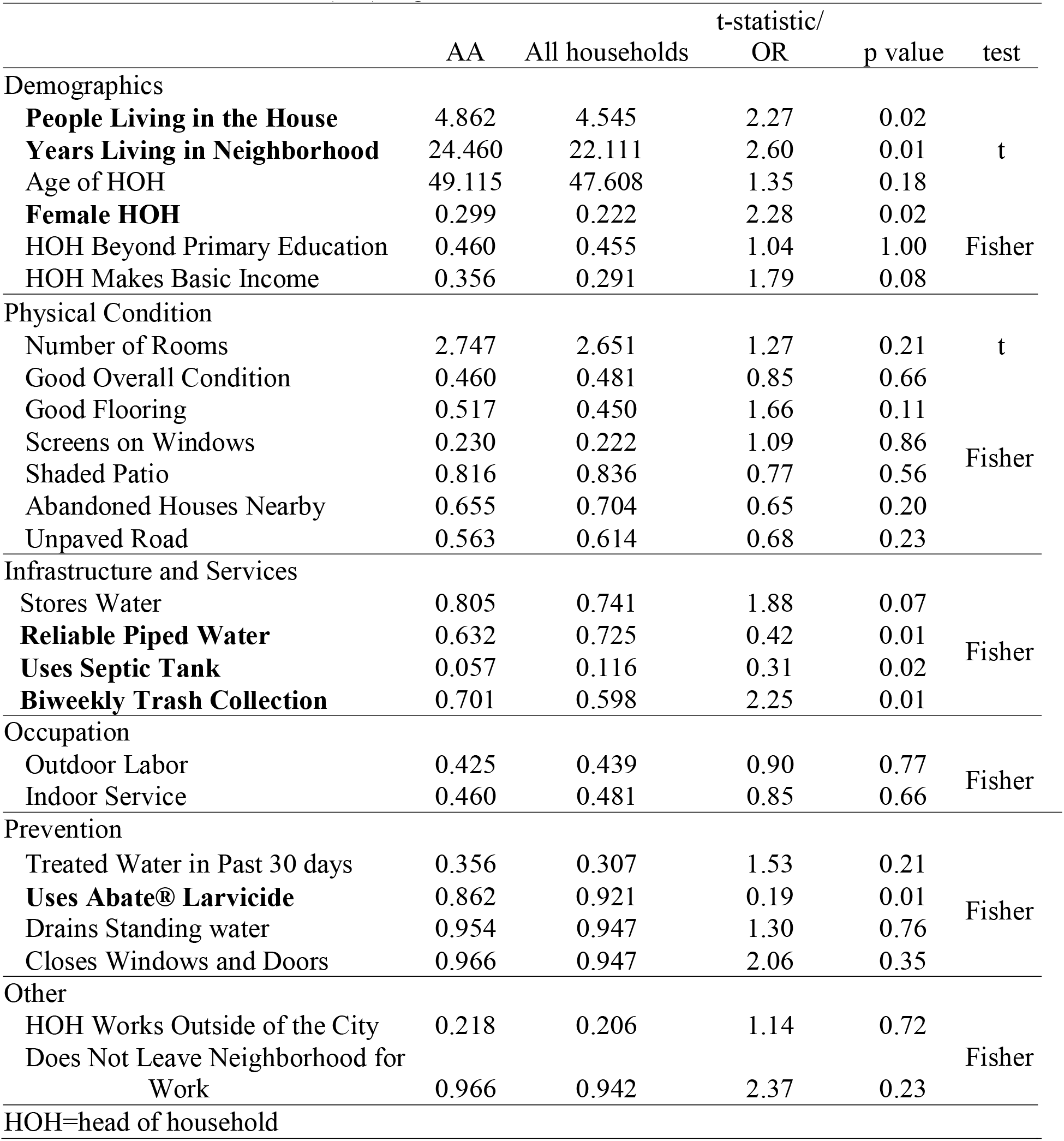
Household level social-ecological factors associated with female *Aedes aegypti* presence (AA) in homes in Huaquillas, Ecuador. Variables given as average value or proportion. Significant associations (p<0.05) are in bold. The t statistic is given when the t test is used and an odds ratio (OR) is given when Fisher’s exact test is used.

### Social-ecological models of household female Aedes aegypti presence

There were eight models within 2 AICc of the top ranked model. All eight models contained the variables: number of years the HOH has lived in the neighborhood (increased exposure), and reliable (un-interrupted) water supply (protective). Additional variables appearing in one of more models, in order of importance (weight) were use of a septic tank, female HOH, the number of people in the household, and biweekly trash collection. Interestingly, the use of larvicide was not found to be important in the top models. The model averaged parameter estimates are given in Table 2, in which odds ratios (OR) denote risk of exposure for OR>1, and protective exposure for OR<1. Relative variable importance among these models is in Table 3. We see that the two most important variables are also the two statistically significant model parameters overall: number of years the HOH has lived in the neighborhood, and reliable water supply.

**Table 2.** SES factors and female *Aedes aegypti* presence in households in Huaquillas, Ecuador. Weighted best-fit model averaged parameter estimates, standard error (SE), odds ratio (OR) and 95% confidence intervals on the ORs included for fixed effects. Bold indicates the variable is statistically significant at α > 0.05. Random effects of cluster and month are given as variance measures, and were included in every model.

| Fixed Effects |  |  |  |  |  |
| --- | --- | --- | --- | --- | --- |
|  | Estimate | SE | Pr(>\|z\|) | OR | 95% CI |
| (Intercept) | - 0.19 | 0.34 | 0.58 | 0.82 | 0.23 - 6.28 |
| **Years Living in Neighborhood** | 0.04 | 0.02 | 0.03 | 1.04 | 1.00 - 1.08 |
| **Reliable Piped Water** | -0.99 | 0.46 | 0.03 | 0.37 | 0.15 – 0.93 |
| Uses Septic Tank | -0.61 | 0.99 | 0.54 | 0.54 | 0.03 - 1.63 |
| People Living in the House | 0.05 | 0.09 | 0.60 | 1.05 | 0.93 - 1.44 |
| HOH Female | 0.11 | 0.32 | 0.73 | 1.12 | 0.60 - 4.82 |
| Biweekly Trash Collection | 0.11 | 0.34 | 0.75 | 1.12 | 0.58 – 5.90 |
| Random Effects |  |  |  |  |  |
| Groups | Variance | Std. Dev |  |  |  |
| Cluster | 0.60 | 0.78 |  |  |  |
| Month | 0.19 | 0.44 |  |  |  |

**Table 3.** Relative importance of each variable included in models. Calculated as a sum of the Akaike weights over all the models in which the variable appears. The number of top models which include the variable is also included.

|  | Summed<br>Akaike Weights | # of Models<br>Containing |
| --- | --- | --- |
| Years Living in Neighborhood | 1.00 | 8 |
| Reliable Water Supply | 1.00 | 8 |
| Uses Septic Tank | 0.40 | 3 |
| People Living in the House | 0.34 | 3 |
| Female HOH | 0.20 | 2 |
| Biweekly Trash Collection | 0.18 | 2 |

## Discussion

In this study we investigated the drivers of household-level female *Ae. aegypti* presence in Huaquillas, Ecuador, to identify social-ecological conditions that promote potential arboviral disease risk to inform vector control and intervention strategies. Precipitation during the study period was anomalously low compared to long-term averages (Fig 1). In several recent studies, the role of drought in altering the way water storage occurs in urban landscapes has been highlighted as a potential key factor in *Ae. aegypti* habitat in urban environments [54–60]. Given this was a particularly dry year, in an already arid environment, the role of precipitation in the timing of *Ae. aegypti* presence may be different than in an average year. We found that the prior week’s precipitation was an important predictor of *Ae aegypti* presence, in combination with the current week’s temperature. Whether the role of precipitation is emphasized or diminished in a dry year is likely mediated by human-driven water storage and use on the landscape. In outdoor, rain-filled habitats, accumulated precipitation can generate oviposition sites for *Ae. aegypti*, but extreme precipitation events can flush out those same larval habitats [61,62]. Thus, the relationship between precipitation and vector population size is not linear, and may depend more on the intensity of precipitation events [9,63]. In our study, *Ae. aegypti* presence was significantly correlated with precipitation lagged by 3 weeks, and 1 week, alone, but when included in a model with temperature, the longer lag dropped out of model importance, and the 1-week lag was found not to be significant, suggesting a stronger role of temperature in this location. In rain-filled habitats, precipitation events increase the suitability of larval habitats, prompting eggs of *Ae. aegypti* to hatch and begin development [64]. The 1-week lag in precipitation likely indicates sufficient humidity and moisture for mosquito activity in the current week, and perhaps trigger egg hatching, but is likely too short a time-frame for development to flying adults. The 3-week lag identified in this study is longer than the typical development time for *Ae. aegypti* [65], however in an arid environment such as this, larval habitat may dry after precipitation events, increasing the time necessary to develop [66,67]. In the urban environment, the timing and degree to which precipitation influences vectors is also highly modulated by the social-ecological environment [11,68]. The role of precipitation may be more identifiable when containers and buckets are visibly on household premises, but is diminished when alternative oviposition sites such as water storage tanks, cisterns, and other water infrastructure sites are available [54,69].

During the study period, mean temperatures in Huaquillas were within historical ranges (Fig 1). Laboratory studies of *Ae. aegypti* have found nonlinear relationships between mean temperature and physiological traits [70,71]. The weekly mean temperatures assessed in our climate analysis range from approximately 26 and 28°C. Biting, development, fecundity, and mortality are positively correlated with mean temperatures within this range [72]. We found that weekly *Ae aegypti* presence was significantly associated with mean temperatures of the same week. This captures the immediate effect of temperature on biting by *Ae. aegypti* in households where people reside [73]. Outdoor temperatures higher than 21°C may drive *Ae. aegypti* indoors to reduce mortality [74], and indoor resting behavior is characteristic of *Ae. aegypti*, especially while processing blood meals [75], so sheltering in shade indoors may be an adaptive strategy for cooling. Prior studies of *Ae. aegypti* in desert climates have occurred in Texas, Arizona, and parts of Mexico [76,77]. In Mexico, a study found differences in the age structure of *Ae. aegypti* populations between two cities in desert and steppe climates, with older *Ae. aegypti* populations in the desert [76]. While precipitation did not differ a lot between the two sites, the cooler steppe population underwent a period of low humidity, which may impact survival. This is important for disease transmission, as *Ae. aegypti* must live long enough to feed and become infectious, and the authors suggested that the higher population turnover in the steppe may contribute to the surprising lack of dengue establishment. This points to the complexities of interactions between temperature (which can exceed optima for survival), precipitation, and sufficient humidity in an arid environment.

In this study, the presence of *Ae aegypti* at the household level differed across months. Changes in *Ae. aegypti* presence at coarse temporal scales are driven by climatic factors [78,79], and increased household *Ae. aegypti* presence in Huaquillas during the study period can be attributed to differences in precipitation and temperature. The seasonal nature of Huaquillas’ climate may thus point to time periods when targeted vector control interventions would be optimally effective.

The household clusters in this study served as a neighborhood-scale measure of space. Presence of *Ae. aegypti* varied significantly between the clusters, meaning there is a high degree of sub-city level spatial variability. Unfortunately, at this sample size, more detailed spatial trends are hard to identify, but point to the need for future studies into heterogeneity at sub-city scale. *Ae. aegypti* has a short dispersal range of about 250 m [35], and its dispersal can be limited by barriers such as major roads or stretches of land unsuitable for oviposition [80]. Several studies have identified distinct vector subpopulations existing in close proximity in urban settings [81–83]. The implications of this in a busy border city call for further studies in this location.

### Social-ecological drivers of risk

In single variable models of *Ae aegypti* presence, the number of years the HOH lived in the neighborhood, the number of people living in the household, and whether the HOH was female were all important and significant demographic components of the social-ecological system. Infrastructurally, reliable piped water supply and trash collection were important and significant, as was whether the household was on a septic tank system, and if the household used Abate^®^ larvicide. Households with *Ae. aegypti* were significantly more likely to use larvicide, which suggests that household members are sensitive to *Ae. aegypti* presence, and may increase preventative actions when the mosquito is detected. This may also mean that control efforts by the Ministry of Health are well targeted at households with *Ae. aegypti*, but may require increased capacity and frequency.

In single variable models, septic tanks were found to be protective against mosquito presence, even though septic tanks act as persistent oviposition sites in other locations [84–86]. In the best-fit model, the role of septic tanks was not significant, so the direction of exposure association (protective or risky) was not clear. Septic tanks in Huaquillas are typically underground with no suitable entrance for mosquitoes (Fig 4), and are used widely in the periphery of the city where sewer infrastructure has only recently become available. All households with septic tanks were in cluster 7 which is in a low-density residential area on the city outskirts. Given that septic tanks occur in peripheral areas where municipal water and sewer infrastructure is a recent addition, and appeared to confer a protective association, this relationship warrants further examination.

While septic tanks, along with female HOH, the number of people living in the household, and trash collection were all important to the best fit model to the social-ecological system model, the two significant variables, the number of years the HOH lived in the neighborhood, and reliability of water supply were the most important and significant social-ecological variables associated with risk of *Ae aegypti* presence at the household level. Interestingly, there was an increase in exposure risk with the number of years HOH lived in the neighborhood, and a protective effect of having reliable water supply. The role of water infrastructure in exposure risk to *Ae aegypti* at the household level has been found in previous studies, and points to the fundamental and vital role reliable water supply and urban infrastructure play in *Ae aegypti* endemic environments. Given the importance of septic tank usage, and the clear role of water supply, this study suggests that urban infrastructure around water supply and use is playing a large role in the risk of *Ae aegypti* presence in the household in Huaquillas.

**Fig 4.**
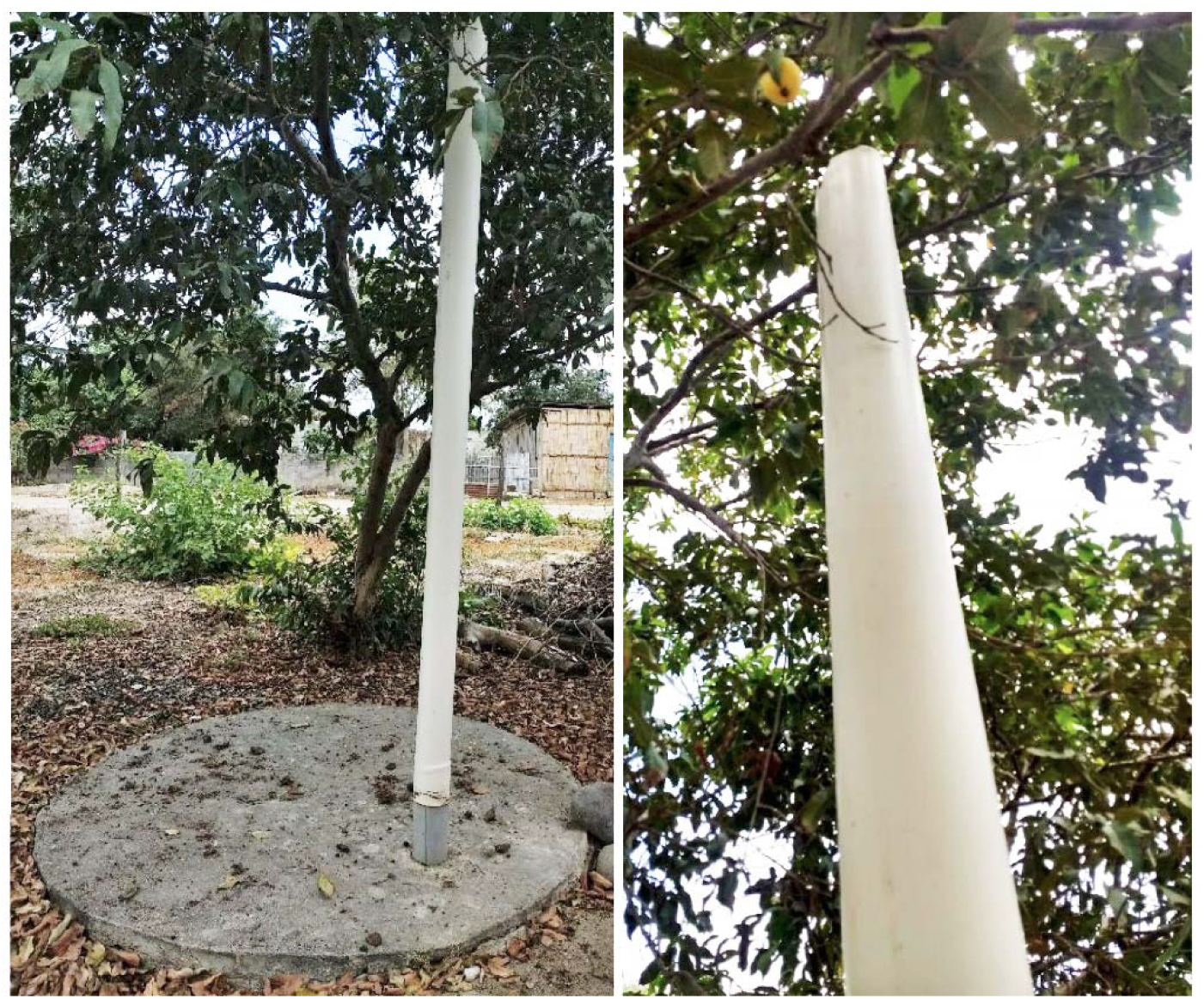
Example of a septic system in Huaquillas, August 2019. Septic systems are commonly used by households not connected to the municipal sewer network. A vent pipe connected to the underground tank rises through a sealed opening in the access cover. The vent pipe stands about 3 meters high. (Photo Credit: R. Sippy)

## Conclusion

In this study we explored climatic and social-ecological factors associated with household-level female *Ae. aegypti* presence, and temporal and spatial trends across an arid border city in Ecuador. The results of our analyses may inform potential control strategies (timing) and interventions (improved water infrastructure) to reduce vector-borne disease risk in the city of Huaquillas in southern coastal Ecuador. Given that this study was conducted in an exceptionally dry year and the evidence for water supply and usage as major factors in household-level risk, water-related interventions at multiple scales could be important.

The social-ecological environment that influences the urban *Ae. aegypti* mosquito varies substantially from place to place. Local studies are especially needed to guide policy and inform interventions. Integrated vector control requires information collection, assessment, and decision making at local scales. While there is value in national to global studies of *Ae. aegypti* populations, these studies produce information that is of most relevance for national or international decision-makers. We acknowledge that there are logistical and resource related challenges inherent in conducting investigations at smaller scales, and in this study, while leveraging an immensely rich dataset over several months, we still ran into issues of small sample size. However, in order to integrate social-ecological systems approaches into essential local-scale work, we suggest that the design and methodological approach of this study is one example of how some of these challenges can be met. As many parts of the world become increasingly urban and ever more connected to global transportation networks, the number of places with endemic *Ae. aegypti* populations will increase. Climate change throughout the 21st century is also set to increase the area suitable for *Ae. aegypti* presence [87]. These developments will further increase the importance of research at multiple scales to guide management and policy. Vector control will continue to be a critical component of arboviral disease prevention, even as additional intervention options become available. Understanding the systems which allow vectors to exist, persist, and transmit disease will remain critical in promoting human health and wellbeing for the foreseeable future.

## Acknowledgments

This study was funded by NSF EEID DEB 1918681 to SJR, EAM, AMS. EAM was also supported by NIH 1R35GM133439-01, the Terman Award, and the Helman Faculty Fellowship.

We thank SUNY Upstate Medical University and the Salud Comunitaria field team for supervising and conducting the data collection necessary for this study. We are also grateful to our collaborators at the Ministry of Health and for all community members who volunteered to participate in this study.

## Supporting information

**S1 Table.**
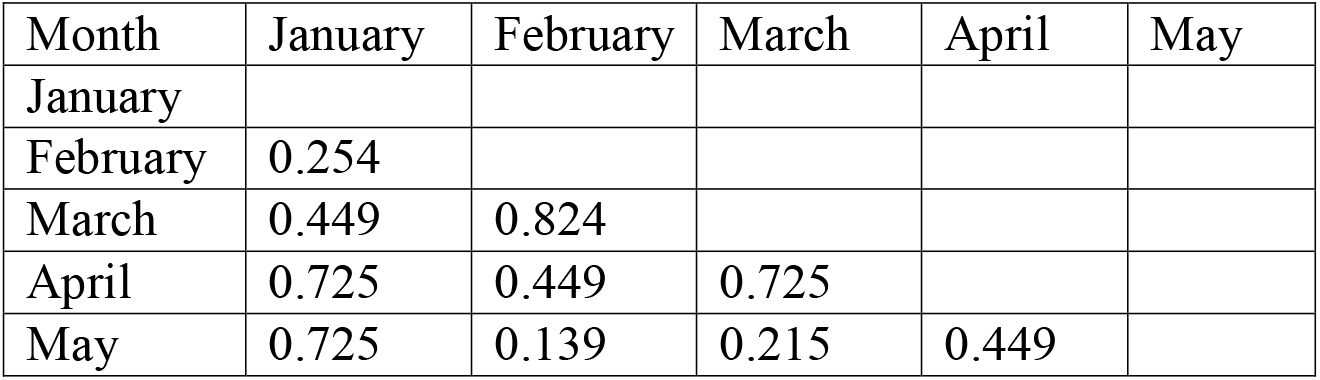
Matrix of *p* values from post hoc tests on pairs of months.

| Month | January | February | March | April | May |
| --- | --- | --- | --- | --- | --- |
| January |  |  |  |  |  |
| February | 0.254 |  |  |  |  |
| March | 0.449 | 0.824 |  |  |  |
| April | 0.725 | 0.449 | 0.725 |  |  |
| May | 0.725 | 0.139 | 0.215 | 0.449 |  |

**S2 Table.**
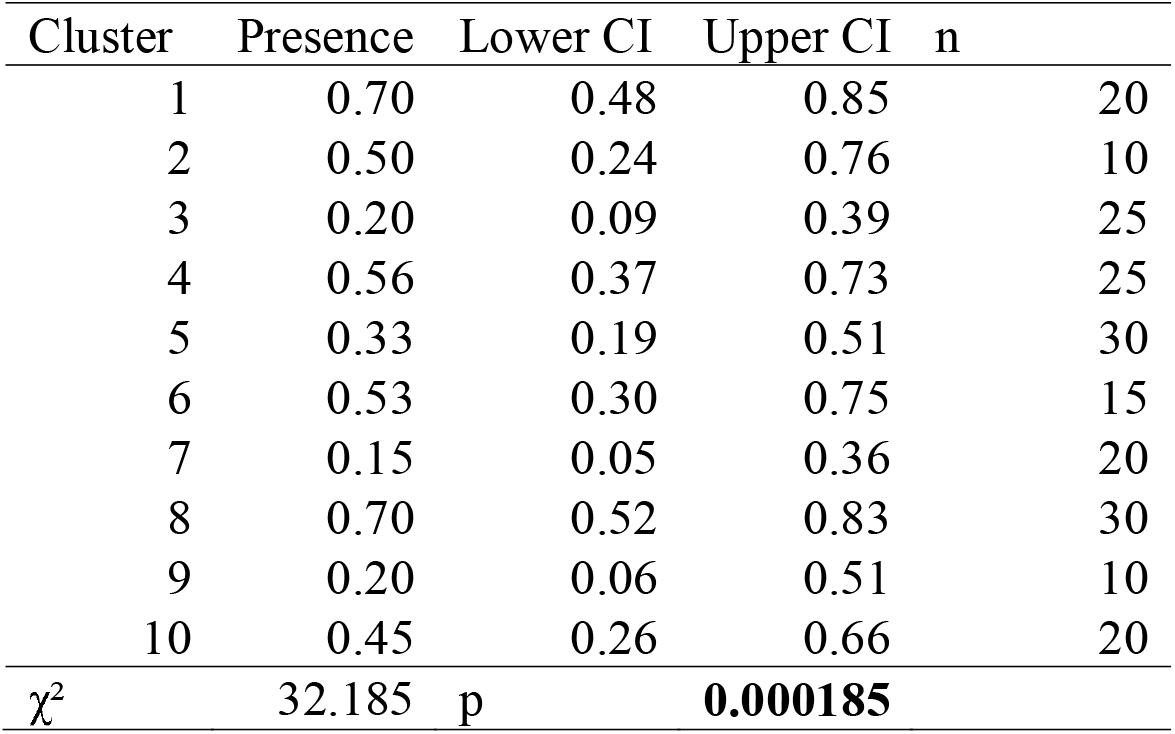
Proportion of households found present for *Aedes aegypti* by cluster.

**S3 Table.** Matrix of p values from post hoc tests on pairs of clusters.

| Cluster | 1 | 2 | 3 | 4 | 5 | 6 | 7 | 8 | 9 | 10 |
| --- | --- | --- | --- | --- | --- | --- | --- | --- | --- | --- |
| 1 |  |  |  |  |  |  |  |  |  |  |
| 2 | 0.617 |  |  |  |  |  |  |  |  |  |
| 3 | **0.012** | 0.258 |  |  |  |  |  |  |  |  |
| 4 | 0.578 | 1.000 | 0.090 |  |  |  |  |  |  |  |
| 5 | 0.090 | 0.642 | 0.578 | 0.258 |  |  |  |  |  |  |
| 6 | 0.656 | 1.000 | 0.154 | 1.000 | 0.426 |  |  |  |  |  |
| 7 | **0.012** | 0.247 | 0.870 | 0.056 | 0.426 | 0.111 |  |  |  |  |
| 8 | 1.000 | 0.500 | **0.008** | 0.599 | 0.060 | 0.573 | **0.007** |  |  |  |
| 9 | 0.090 | 0.578 | 1.000 | 0.246 | 0.866 | 0.426 | 1.000 | 0.060 |  |  |
| 10 | 0.426 | 1.000 | 0.258 | 0.713 | 0.713 | 0.874 | 0.247 | 0.250 | 0.462 |  |

